# Arf6 Regulates Endocytosis and Angiogenesis by Promoting Filamentous Actin Assembly

**DOI:** 10.1101/2023.02.22.529543

**Authors:** Caitlin R. Francis, Makenzie L. Bell, Marina M. Skripnichuk, Erich J. Kushner

**Affiliations:** Department of Biological Sciences, University of Denver, Denver, CO

**Keywords:** Arf6, Actin, Clathrin-Mediated Endocytosis, Angiogenesis, Lumenogenesis, Blood vessel Development, ARNO, ACAP2, GEF, GAP

## Abstract

Clathrin-mediated endocytosis (CME) is a process vital to angiogenesis as well as general vascular homeostasis. In pathologies where supraphysiological growth factor signaling underlies disease etiology, such as in diabetic retinopathy and solid tumors, strategies to limit chronic growth factor signaling by way of CME have been shown to have tremendous clinical value. ADP ribosylation factor 6 (Arf6) is a small GTPase that promotes the assembly of actin necessary for CME. In its absence, growth factor signaling is greatly diminished, which has been shown to ameliorate pathological signaling input in diseased vasculature. However, it is less clear if there are bystander effects related to loss of Arf6 on angiogenic behaviors. Our goal was to provide a analysis of Arf6’s function in angiogenic endothelium, focusing on its role in lumenogenesis as well as its relation to actin and CME. We found that Arf6 localized to both filamentous actin and sites of CME in 2-dimensional culture. Loss of Arf6 distorted both apicobasal polarity and reduced the total cellular filamentous actin content, and this may be the primary driver underlying gross dysmorphogenesis during angiogenic sprouting in its absence. Our findings highlight that endothelial Arf6 is a potent mediator of both actin regulation and CME.

## INTRODUCTION

Endothelial cells require tight control of endocytic processes to integrate both intrinsic and extrinsic signaling into large-scale morphogenic events during blood vessel formation. This is best illustrated by the dependency of critical receptors such as vascular endothelial growth factor receptor 2 (VEGFR2), or other receptor tyrosine kinase family members, on endocytic processes for activation[1–4]. Specifically, internalization of these receptors propagates a signaling response as well as timely quenching of receptor activity[5]. This removal of proteins from the plasma membrane is largely carried out by a well-characterized process termed clathrin-mediated endocytosis (CME)[6–8]. Disruption in CME has been shown to drastically affect gross blood vessel development *in vitro* and *in vivo* as CME plays a fundamental role in a vast number of critical cellular processes[9, 10]. Endocytosis by way of CME is not unique to endothelial tissue; however, the accessory proteins that adapt CME to biological scenarios exclusive to blood vessel morphogenesis have yet to be fully explored.

ADP-ribosylation factor (Arf) GTPases are a sub-family of small GTPases with six isoforms[11]. In particular, Arf6 has been reported to operate at the plasma membrane with a well-defined function in promoting the assembly of filamentous actin at sites of CME[12]. The predominant model is that Arf6 in its GTP form catalyzed by its guanine exchange factor (GEF) ARNO[13, 14] localizes to sites of clathrin assembly and modifies plasma membrane composition favoring recruitment and activation of Rac1 and ARP2/3 promoting actin assembly[13]. Localized polymerization of actin at clathrin-coated pits provides the scaffolding for motor proteins to generate sufficient pulling force for pit internalization and scission[15]. In the absence of Arf6 or expression of a dominant-negative mutant, generalized CME programs are distorted[16]. Adding to the complexity of Arf6’s biological function is evidence for non-CME related roles of Arf6. For instance, Arf6 does not only localize to sites of CME, but is also colocalized to areas of high-actin density, suggesting a regulatory role in cell motility-related actin assembly by interfacing with other GTPases such as Rab35[17].

Arf6’s function in blood vessel development has been investigated primarily through its requirement in VEGFR2 signaling and relation to cancer progression[18–21]. Endothelial-specific ablation of Arf6 in a mouse model of diabetic retinopathy protected against vascular leakage by reducing VEGFR2 signaling capacity[22]. This investigation and others[23] in endothelial cells demonstrate that Arf6 is required for CME of VEGFR2[24, 25]; thus, the therapeutic potential lies in its ability to reduced VEGF signaling in diseases where chronic VEGFR2 activation, or other growth factor pathways, becomes pathological. As there are multiple ways to specifically target CME, another clinically attractive trait of using Arf6 inhibition is that Arf6 is also vital for clathrin-independent endocytosis[26]. With regard to reducing pathological angiogenesis, Arf6 inhibition may be more advantageous than solely targeting CME as there is strong evidence suggesting VEGFR2 is also regulated through clathrin-independent pathways[4]. Although, this is a somewhat tangential avenue for limiting growth factor signaling, it is clearly efficacious and medically relevant.

Given the utility of Arf6 inhibition as a viable chemotherapeutic since the recent advent of small molecule inhibitors[27], we sought to provide a more holistic understanding of Arf6’s function in endothelial tissue with regard to its localization in angiogenic endothelium, participation in lumen formation behaviors as well as its relation to actin and CME. Our findings both validate and extend Arf6’s function in blood vessel regulation. True to a more promiscuous role, we found that Arf6 localized to both filamentous actin and sites of CME in 2-dimensional (2D) culture. Interestingly, in 3D sprouts, Arf6 strongly localized with apical actin and other luminal proteins. Loss of Arf6 distorted both apicobasal polarity and resident protein amounts in sprouts. Reasoning that the primary defect of Arf6 ablation was related to its influence on actin polymerization, we tested for global cellular shifts in actin pools (e.g. globular vs filamentous). Our results suggest that loss of Arf6 reduced the total cellular filamentous actin content, and this may be the primary driver underlying gross dysmorphogenesis during angiogenic sprouting in its absence. Overall, our findings highlight that endothelial Arf6 is a potent mediator of actin regulation and CME. Although, distorting the Arf6 pathway is capable of attenuating pathological growth factor signaling in various disease models, it also carries potential deleterious effects on vital angiogenic behaviors such as sprouting and lumen formation due to its central role in actin regulation.

## MATERIALS AND METHODS

### Data Availability

The authors will make any data, analytic methods, and study materials available to other researchers upon written request.

### Experimental Procedures

All research complied with the University of Denver Institutional Biosafety Committee (IBC).

### Reagents

All reagents, siRNA and plasmid information are listed in the reagents table in the Supplementary Information (**Supplementary Tables 1-5**).

### Cell Culture

Pooled human umbilical vein ECs cultured in proprietary media (PromoCell Growth Medium, ready-to-use) for 2 to 5 passages. For experiments, glass-bottomed imaging dishes were exposed to deep UV light for 6 minutes and coated with Poly-D-Lysine for a minimum of 20 minutes. Small interfering RNA was introduced into primary human umbilical vein ECs using the Neon transfection system (ThermoFisher). See Supplementary Table 5 for sources of siRNA. All siRNA were resuspended to a 20 μmol/L stock concentration and used at 0.5 μmol/L. Normal human lung fibroblasts and HEK-A were maintained in DMEM supplemented with 10% fetal bovine serum and antibiotics. Both normal human lung fibroblasts and HEKs were used up to 15 passages. All cells were maintained in a humidified incubator at 37 °C and 5% CO_2_.

### Sprouting Angiogenesis Assay

Fibrin-bead assay was performed as reported by Nakatsu et al.[28, 29]. Briefly, human umbilical vein ECs were coated onto microcarrier beads and plated overnight. SiRNA-treatment or viral transduction was performed the same day the beads were coated. The following day, the EC-covered microbeads were embedded in a fibrin matrix. Once a clot was formed, media was overlaid along with approximately 100,000 normal human lung fibroblasts. Media was changed daily along with monitoring of sprout development. Sprout characteristics were quantified in the following manner. Sprout numbers were determined by counting the number of multicellular sprouts (sprouts that did not contain at least 3 cells were not counted) emanating from an individual microcarrier beads across multiple beads in each experiment. Sprout lengths were determined by measuring the length of a multicellular sprout beginning from the tip of the sprout to the microcarrier bead surface across multiple beads. Percent of non-lumenized sprouts were determined by quantifying the proportion of multicellular sprouts whose length (microcarrier bead surface to sprout tip) was <80% lumenized across multiple beads. Sprout widths were determined by measuring the sprout width at the midpoint between the tip and the microcarrier bead across multiple beads. Experimental repeats are defined as an independent experiment in which multiple cultures, containing numerous sprouting beads were quantified; this process of quantifying multiple parameters across many beads and several cultures was replicated on different days for each experimental repeat.

### Lentivirus and Adenovirus Generation and Transduction

Lentivirus was generated by using the LR Gateway Cloning method[30]. Genes of interest and fluorescent proteins were isolated and incorporated into a pME backbone via Gibson reaction[31]. Following confirmation of the plasmid by sequencing the pME entry plasmid was mixed with the destination vector and LR Clonase. The destination vector used in this study was pLenti CMV Neo DEST (705-1) (gift from Eric Campeau & Paul Kaufman; Addgene plasmid #17392). Once validated, the destination plasmids were transfected with the three required viral protein plasmids: pMDLg/pRRE (gift from Didier Trono; Addgene plasmid # 12251), pVSVG (gift from Bob Weinberg; Addgene plasmid #8454) and psPAX2 (gift from Didier Trono; Addgene plasmid #12260) into HEK 293 cells. The transfected HEKs had media changed 4 hours post transfection. Transfected cells incubated for 3-4 days and virus was harvested.

### Membrane Fraction Assay

Membrane fractions were performed according to the guidelines provided using the Thermo-Scientific Mem-PER Plus Membrane Protein Extraction Kit.

### Detection of Globular and Filamentous Actin

Globular and filamentous actin ratios were determined by western blot as described by commercially available G-actin/ F-actin In Vivo Assay Kit (Supplemental Table 1). Globular and filamentous immunocytochemistry was performed as previously described [32]. Briefly, cells were fixed with 4% PFA for 10 minutes and permeabilized in ice cold acetone for 5 minutes and washed. Cells were then incubated for 15 minutes in 2% BSA with globular actin-binding protein GC globulin (Sigma). Following incubation, cells were washed three times in PBS. After washes cells incubated with an anti-GC antibody in BSA for 15 minutes, washed three times, and incubated in anti-rabbit-555 secondary prior to imaging.

### Immunoblotting and Protein Pull-Down

HUVEC cultures were trypsinized and lysed using Ripa buffer (20 mM Tris-HCl [pH 7.5], 150 mM NaCl, 1 mM Na2 EDTA, 1 mM EGTA, 1% NP-40, 1% sodium deoxycholate, 2.5 mM sodium pyrophosphate, 1 mM β-glycerophosphate, 1 mM Na3VO4, 1 μg/mL leupeptin) containing 1× ProBlock^™^ Protease Inhibitor Cocktail-50 (GoldBio) and processed as previously described[17]. Protein was then transferred to Immun-Blot PVDF Membrane at 4°C, 100 V for 1 hour 10 minutes. Blots were blocked in 2% milk proteins for 1 hour, then put in primary antibody at specified concentrations overnight. After 3 10-minute washes with PBS, secondary antibodies at specified concentrations were applied for 4 hours. After 3 additional PBS washes, blots were developed with ECL reagent. Arf6 activation assay blots were performed using commercially available kits listed in the Supplemental Information.

### Immunofluorescence and Microscopy

For immunofluorescence imaging of 2-dimensional cells, prior to seeding cells, coverslips were treated with poly-D Lysine for 20 minutes and washed twice with PBS. HUVECs were fixed with 4% paraformaldehyde (PFA) for 7 min. ECs were then washed three times with PBS and permeabilized with 0.5% Triton-X for 10 minutes. After permeabilization, cells were washed three times with PBS. ECs were then blocked with 2% bovine serum albumin (BSA) for 30 min. Once blocked, primary antibodies were incubated for approximately 4–24 hours. Thereafter, primary antibodies were removed, and the cells were washed 3 times with PBS. Secondary antibody with 2% BSA were added and incubated for approximately 1–2 hours, washed 3 times with PBS, and mounted on a slide for imaging. All primary and secondary antibodies are listed in the Supplemental Information table 3.

For imaging the fibrin-bead assay, fibroblasts were removed from the clot with a 1-minute trypsin incubation. Following incubation, the trypsin was neutralized with DMEM containing 10% BSA, washed three times with PBS, and fixed using 4% paraformaldehyde for 40 minutes. After fixation, the clot was washed three times with PBS, permeabilized with 0.5% Triton-X for 2 hours and then blocked with 2% BSA for 1 hour before overnight incubation with primary antibodies. The following day, primary antibodies were removed, and the clot was washed five times with PBS and secondary antibody was added with 2% BSA and incubated overnight. Before imaging, the clot was washed five times with PBS. All primary and secondary antibodies are listed in the Supplemental Information. Images were captured on a Nikon Eclipse Ti inverted microscope equipped with a CSU-X1 Yokogawa spinning disk field scanning confocal system and a Hamamatsu EM-CCD digital camera. Images were captured using a Nikon Plan Apo 60x NA 1.40 oil objective using Olympus type F immersion oil NA 1.518, Nikon Apo LWD 20× NA 0.95 or Nikon Apo LWD 40× NA 1.15 water objective. All images were processed using ImageJ (FIJI).

### Quantification of Fluorescence Intensity

Fluorescence intensity was determined by first projecting the entire cell or sprout to a single image, setting the pixel scale, and then designating a region of interest. The resulting integrated density measurement was then divided by the area to account for fluctuations in cell/sprout size. For quantifying western blot band intensity between groups, the bounding box was set constant from band to band and fluorescence intensity was compared with equal areas and expressed as a ratio to a loading control protein.

### Statistical Analysis

Experiments were repeated a minimum of three times. Statistical analysis and graphing were performed using GraphPad Prism software. Statistical significance was assessed with a student’s unpaired t-test for a two-group comparison. Multiple group comparisons were carried out using a one-way analysis of variance (ANOVA) followed by a Dunnett multiple comparisons test. Data was scrutinized for normality using Kolmogorov-Smirnov (K-S) test. Statistical significance set a priori at p<0.05.

## RESULTS

### Arf6 Localizes to Cortical Actin and Clathrin in 2D Endothelial Cells and 3D sprouts

Several investigations have shown that Arf6 is vital to VEGFR2 and hepatocyte growth factor receptor CME and subsequent signaling in endothelium[22, 33–35]. Surprisingly, to our knowledge, no reports have mapped where Arf6 is localized in endothelial cells (ECs) and sprouting vessels relative to established membrane domains. To address this, we first live-imaged ECs co-expressing the actin protein mCherry-Arp2 and a wild-type (WT) cyan fluorescent protein (CFP)-tagged Arf6. Here, Arf6 and Arp2 showed strong co-localization at membrane accumulations, presumably actin-based peripheral membrane ruffling (**Figure 1A, Movie 1**). Live imaging of tagRFP-Clathrin and WT Arf6-CFP showed that nascent clathrin puncta were also associated with Arf6 (**Figure 1A, Movie 2**). Interestingly, both proteins did not move in perfect synchrony, rather step movements in clathrin puncta would be followed by a lagged recruitment of Arf6.

**Figure 1.**
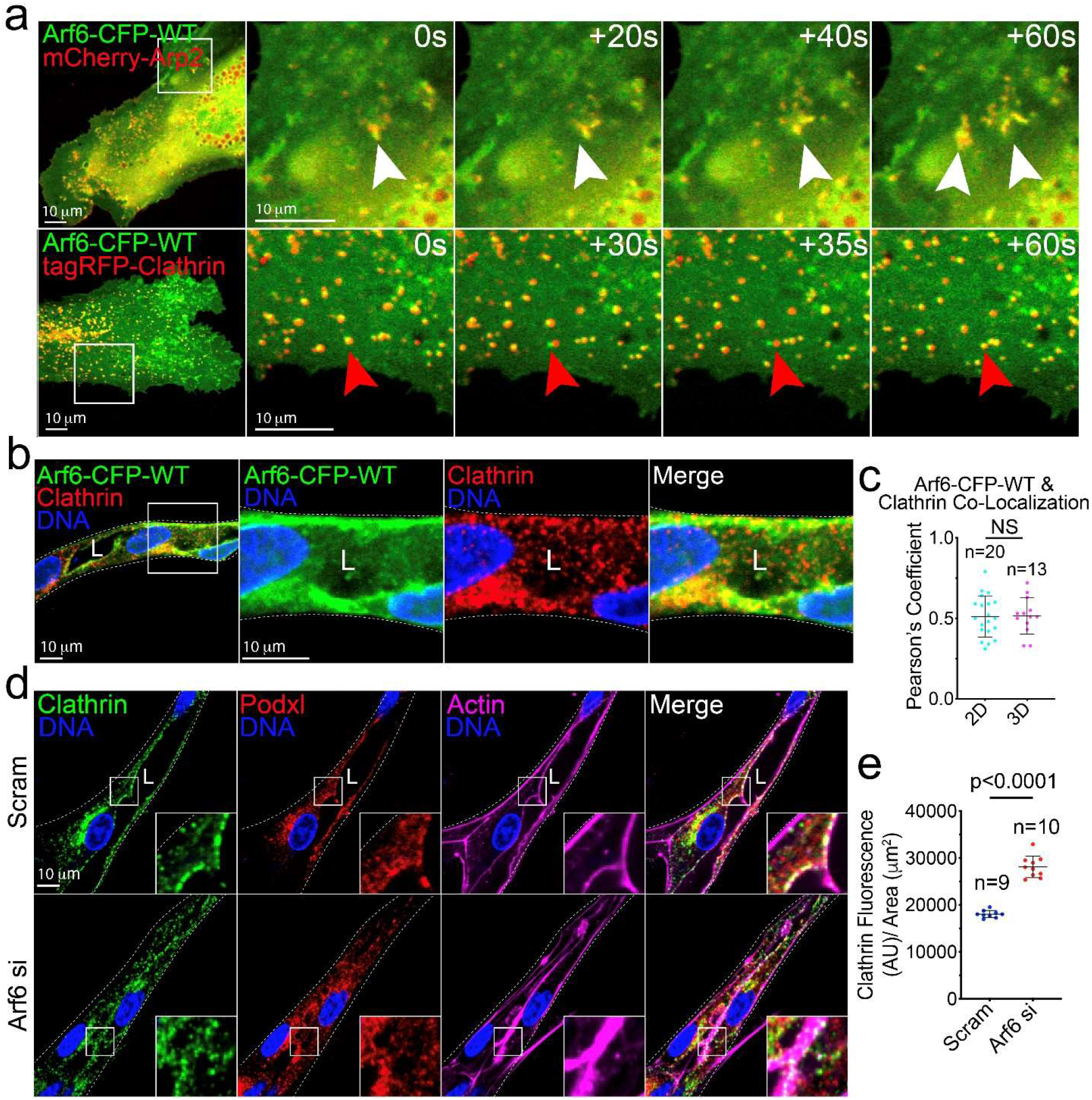
Arf6 localizes to Actin and Clathrin in 2D Culture and 3D sprouts. **A.** Live imaging of wild-type (WT) Arf6-CFP and mCherry-Arp2 (top panels) or tagRFP-clathrin (bottom panels) over indicated timepoints. White arrowheads denote co-localization between Arf6 and Arp2. Red arrowheads denote movement of Arf6 following clathrin. **B.** Image representative of WT Arf6-CFP in sprout structures stained for endogenous clathrin. **C.** Pearson’s Coefficient between Arf6-CFP and clathrin in 2D culture and in sprout structures. **D.** Representative image of scramble (Scram) and Arf6 siRNA (si) knockdown (KD) sprouts stained for clathrin, podocalyxin (Podxl) and actin. **E.** Quantification of clathrin fluorescence intensity for indicated conditions. AU is arbitrary unit. In all panels L denotes lumen. Statistical significance was assessed with an unpaired t-test or a 1-way ANOVA followed by a Dunnett multiple comparisons test. Insets are areas of higher magnification. Error bars represent standard deviation, middle bars represent the mean. White dashed lines mark sprout exterior. All experiments were done using Human umbilical vein endothelial cells in triplicate.

Next, we sought to validate the above association in endothelial sprouting structures as 2D culture affects protein localization due to the lack of an established apicobasal polarity[36]. To test this, we employed a fibrin-bead assay in which ECs sprout from a microcarrier bead and reliably reproduce normal blood vessel sprouting, branching and lumenization behaviors[37, 38]. Using this method, we transduced sprouts with a WT Arf6-CFP virus and stained for endogenous clathrin (**Figure 1B)**. We observed that Arf6 was located at clathrin accumulations to the same degree as non-polarized 2D culture (**Figure 1C**). This data suggests that the presence of an established apicobasal domain does not affect Arf6’s recruitment to clathrin depots. To validate a dependency of CME on Arf6, we knocked down (KD) Arf6 and compared the relative amount of clathrin intensity between groups as a proxy for the amount of active CME sites. Loss of Arf6 significantly increased the amount of clathrin in sprouts as compared with controls (**Figure 1D,E**). Morphologically, clathrin was in close proximity to cortical actin with and without Arf6 KD in sprouts (**Figure 1D**).

Next, we determined where Arf6-CFP localized with regard to filamentous (F)-actin, cortical actin (mCherry-Arp2), or sites of CME (tagRFP-clathrin). We observed significantly greater co-localization of Arf6-CFP with mCherry-Arp2 and tagRFP-Clathrin compared with cytosolic tdTomato (control) or LifeAct (**Supplemental Figure 1A,B**). This finding suggests that Arf6 is preferentially recruited to sites of active actin polymerization. We also co-expressed the constitutively-active or dominant-negative Arf6 mutant with Arp2 or clathrin. Our results again show that active Arf6 is most strongly associated with branch actin marked by Arp2 and clathrin pits (**Supplemental Figure 1C-E**). These results indicate that Arf6 equally localizes to both cortical actin as well as clathrin-associated pits in 2D culture and multicellular sprouts. Generally, these observations are consistent with previous reports in non-endothelial systems showing Arf6’s preference for actin and sites of CME.

### Arf6 is Required For Maintenance of Trans-Membrane Protein Turnover

Unlike 2D culture, sprouts possess an intrinsic apicobasal axis that could influence Arf6 localization. As such, we determined if Arf6 demonstrated localization preference relative to various established apical and basal membrane markers. In aggregate, Arf6 primarily localized to the plasma membrane (**Supplemental Figure 2A; Movie 3**). More specifically, we observed that Arf6 strongly colocalized with the apical protein podocalyxin and phosphorylated-Tie2 (**Figure 2A,B**). Endogenous VEGFR2 puncta and fluorescently-tagged Arf6 demonstrated strong colocalization (**Figure 2A,B**). Lastly, Arf6 has been suggested to have a role in integrin CME and recycling primarily in 2D culture in non-endothelial tissues[39]. In polarized sprouts, Arf6 and β1-integrin did show significant colocalization; although, Arf6 was primarily localized on the apical membrane opposite β1-integrin on the basal surface (**Figure 2A,B**). These results suggest that Arf6 is largely apically localized, perhaps due to its association with resident cortical actin in these areas.

**Figure 2.**
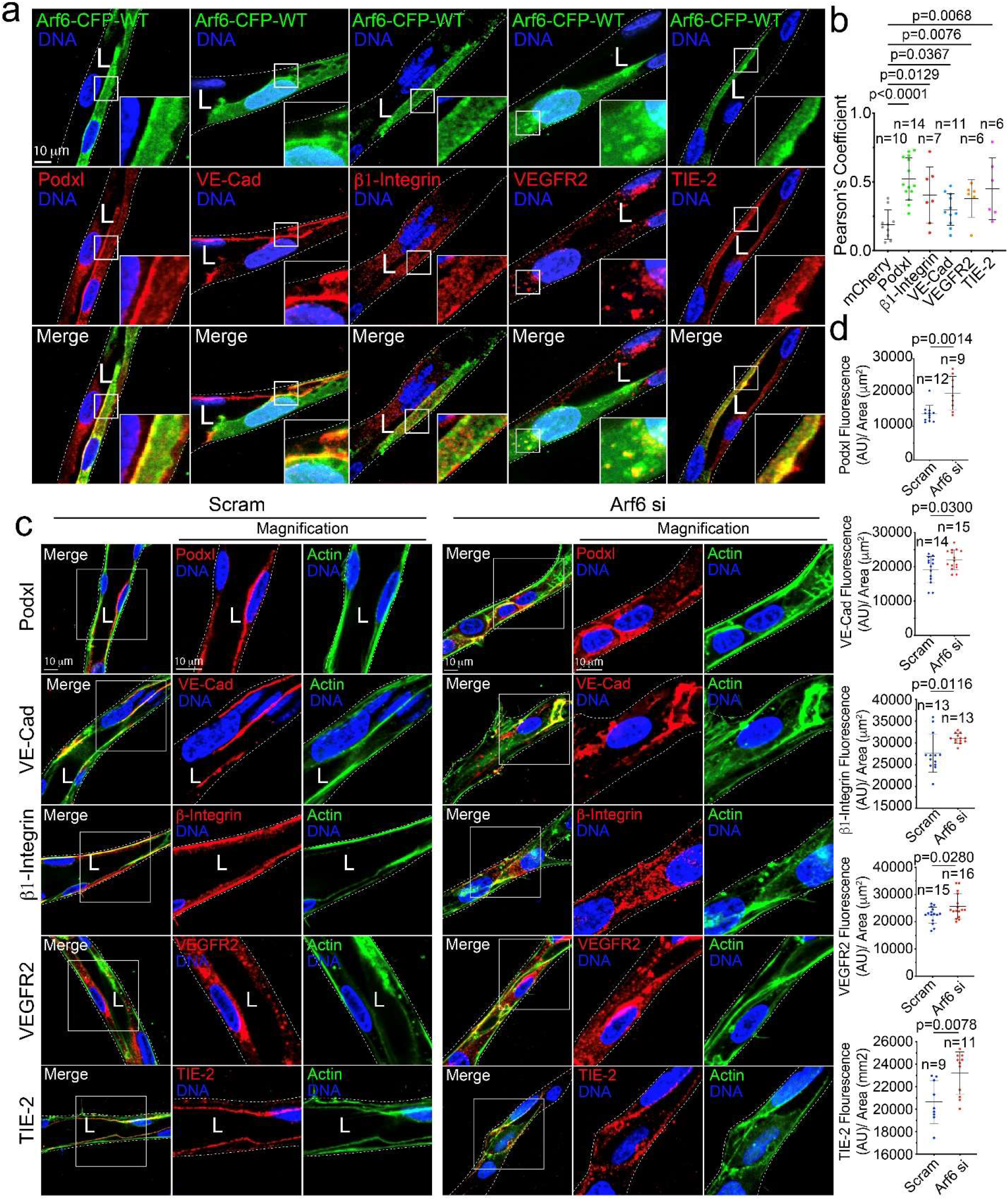
Arf6 is Required for Trans-membrane Localization and Internalization. **A.** Wild-type (WT) Arf6-CFP localization relative to podocalyxin (Podxl), VE-Cadherin (VE-Cad), β1-Integrin, Vascular Endothelial Growth Factor Receptor 2 (VEGFR2), and TIE-2. **B.** Pearson’s Coefficient between Arf6-CFP and indicated proteins. **C.** Representative images of scramble (Scram) control and Arf6 siRNA (si) knockdown (KD) sprouts stained for Podxl, VE-Cad, β-integrin, VEGFR2, and TIE-2. Actin (green) delineates sprout morphology. **D.** Quantification of fluorescence intensity for indicated proteins. In all panels n = number of sprouts. AU is arbitrary unit. Statistical significance was assessed with an unpaired t-test or a 1-way ANOVA followed by a Dunnett multiple comparisons test. Insets are areas of higher magnification. Error bars represent standard deviation, middle bars represent the mean. L denotes lumen. White box is area of magnification. Insets are areas of higher magnification. Dashed lines mark sprout exterior. All experiments were done using Human umbilical vein endothelial cells in triplicate.

As CME and by extension Arf6 are involved in a multitude of endocytic events, we next tested how critical membrane-bound proteins were then affected by loss of Arf6. First, we KD Arf6 and quantified the presence of clathrin puncta as a proxy for the number of CME sites. Loss of Arf6 significantly increased the amount of clathrin (**Figure 1E**) as well as puncta lifetime as compared with controls (**Supplemental Figure 2B**), suggesting in the absence of Arf6 CME can be initiated. To investigate the hypothesis that loss of Arf6 stalls or disrupts CME in angiogenic endothelium, we compared the relative amounts and localization of several apical and basal proteins in sprouts. In terms of protein localization, all assayed proteins demonstrated dysmorphic spatial organization in sprouts compared with control ECs when Arf6 was depleted (**Figure 2C**). Knockdown of Arf6 significantly increased the cellular content of all proteins, indicating that said proteins exocytic trafficking were normal, but are then essentially trapped on the plasma membrane as CME was dysfunctional in the absence of Arf6 (**Figure 2C,D**).

To further confirm this notion, we chemically isolated the plasma membrane and compared amounts of membrane-bound proteins with and without Arf6 depletion (**Figure 3A,B**). Podocalyxin and VE-cadherin demonstrated a significant increase membrane retainment in the absence of Arf6, while β1-integrin, VEGFR2 and Tie-2 demonstrated normal levels. Given both β1-integrin and VEGFR2 were previously shown to be affected by Arf6, we employed a more sensitive method using an antibody feeding assay to quantify the membrane-bound to internalized protein populations. Using this assay as a marker for endocytic capacity, we tracked the ability of ECs to internalize the aforementioned proteins overtime as compared to a 4°C cold-blockade negative control. Knockdown of Arf6 significantly reduced the endocytic capacity of ECs to internalize β1-integrin and VEGFR2 compared to a scramble-treated control (**Figure 3C-E)**. Overall, this finding aligns with the idea that Arf6 plays a fundamental role in CME in which loss of Arf6 results in halted protein internalization, presumably for any protein that employs CME as its chief mechanism of internalization.

**Figure 3.**
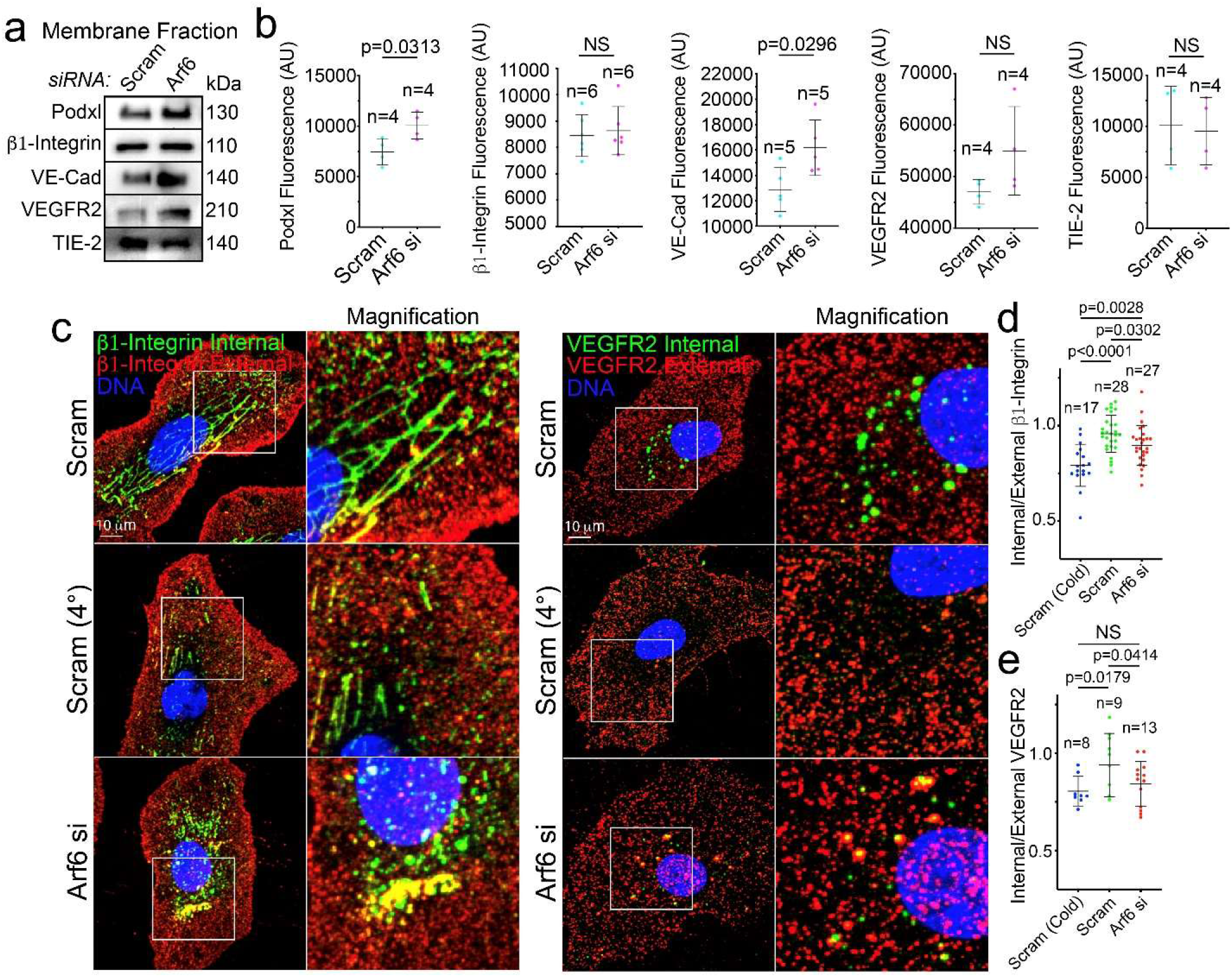
Arf6 is an Indiscriminate Endocytic Regulator. **A.** Western blot of membrane isolations treated with scramble (Scram) or Arf6 siRNA (si). **B.** Quantification of band intensity in membrane fractions in panel A. n = individual membrane fractionation experiments. **C.** Antibody feeding assay representative images differentially stained proteins between siRNA-treated groups. Green channel represents internalized protein and red channel represents external protein. **D-E.** Ratio of internal to external protein. n = number of cells. Error bars represent standard deviation, middle bars represent the mean. Statistical significance was assessed with an unpaired t-test or a 1-way ANOVA followed by a Dunnett multiple comparisons test. NS = Not Significant. All experiments were done using human umbilical vein endothelial cells in triplicate.

### Arf6 Promotes the Assembly of Actin

Since endothelial Arf6 was not only at sites of CME, but was also heavily localized to peripheral actin in 2D culture and cortical actin in sprouts, we next tested how Arf6 influenced cellular actin dynamics. First, we imaged live-cell actin dynamics in 3D sprouts. In Arf6 KD ECs, we observed a thinner network of filaments with an abundance of small actin accumulations leading to a generally disorganized appearance in the actin architecture compared with controls (**Figure 4A, Supplemental Movie 4**). Quantification of F-actin intensity was significantly lower in Arf6 KD sprouts compared with controls, a finding consistent with Arf6 mediating actin polymerization (**Figure 4B; Supplemental Movie 5**). To confirm that Arf6 promoted actin polymerization, we compared the amounts of globular (G), or monomeric, actin to F-actin between groups by differential centrifugation. F-actin was significantly reduced in Arf6 KD ECs as compared with controls (**Figure 4C,D**). Similarly, we stained for G-actin and F-actin in ECs and compared the relative intensities. Again, the ratio of G-to F-actin was elevated in the in Arf6 KD ECs indicating that loss of Arf6 is associated with reduced F-actin (**Figure 4E,F**). These data indicate that loss of Arf6 can greatly affect cellular actin dynamics.

**Figure 4.**
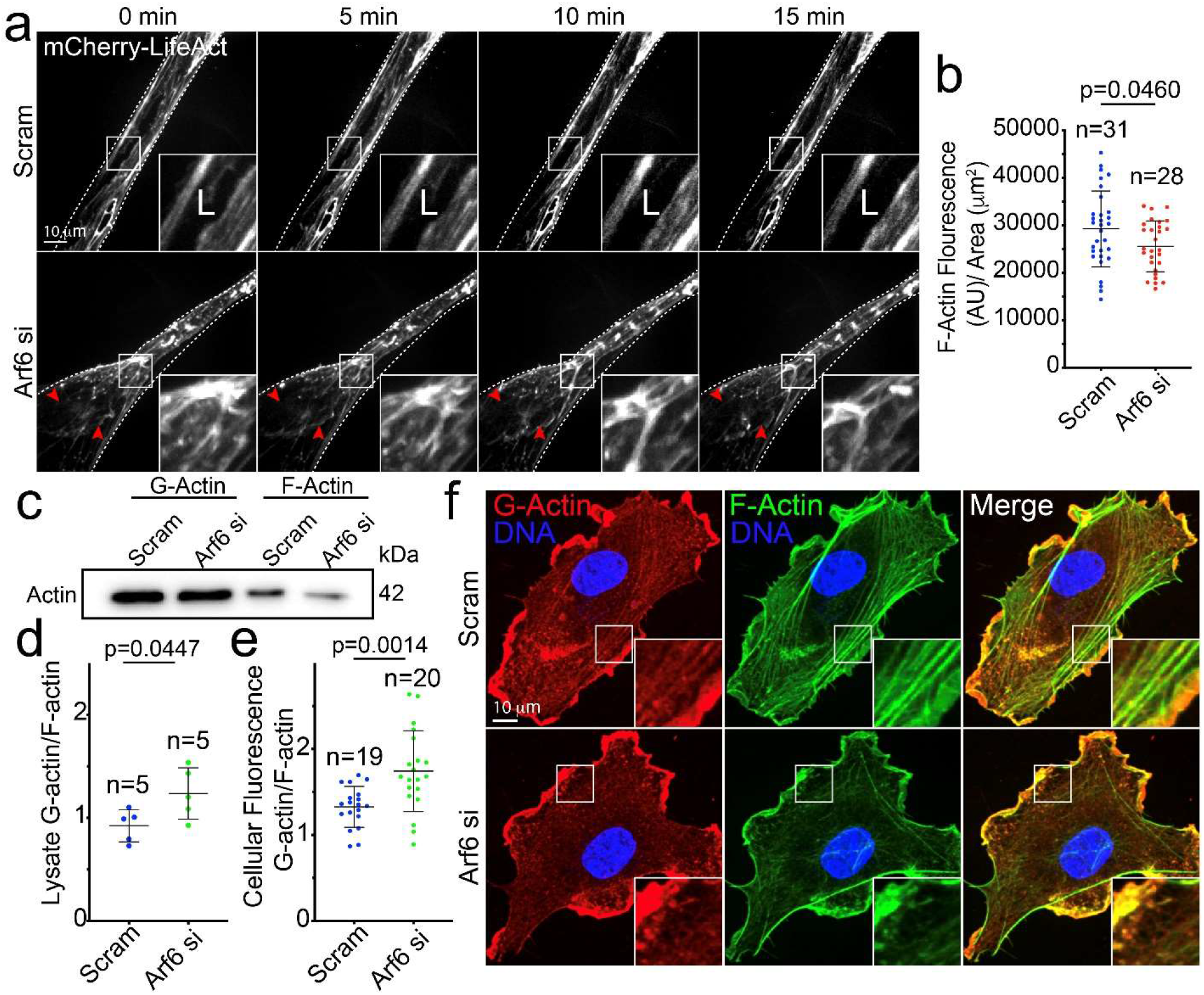
Arf6 Promotes Actin Assembly. **A.** Live imaging of scramble (Scram) control and Arf6 siRNA (si) knockdown (KD) sprouts expressing mCherry-LifeAct lentivirus at indicated timepoints. Red arrowheads denote sites of diminished filamentous actin. Dashed line denotes sprout exterior. L denotes lumen. **B.** Quantification of filamentous-actin (F-Actin) fluorescence intensity in Scram and Arf6 si-treated sprouts. n = number of sprouts. AU is arbitrary unit. **C.** Western blot of globular (G) and filamentous (F) actin in indicated groups. **D.** Quantification of the ratio of globular to filamentous actin from blots represented in panel (C). n = number of blots. **E.** Quantification of the ratio of globular to filamentous actin fluorescence intensities. n = number of cells. **F.** Representative images of cells stained for globular (G-Actin) and F-Actin between indicated conditions. In all images white box denotes area of magnification. NS = non-significant. Error bars represent standard deviation, middle bars represent the mean. Statistical significance was assessed with an unpaired t-test or a 1-way ANOVA followed by a Dunnett multiple comparisons test. All experiments were done in Human umbilical vein endothelial cells in triplicate.

We were intrigued by the results implicating Arf6 as a potent regulator of actin polymerization in ECs. We questioned to what extent the loss of actin polymerization ability, per se, would affect endocytosis. In other words, could we phenocopy the Arf6 KD effect on protein accumulation by simply inhibiting actin polymerization? To test this, we treated ECs with the Arp2/3 inhibitor CK-666[40] to block the formation of branched actin and then quantified the relative protein amounts between conditions. Application of CK-666 significantly increased VE-cadherin, phosphorylated Tie2 and β 1-integrin amounts as compared with controls in 2D culture. This data suggests that having the ability to polymerize branched actin is necessary for protein internalization from the plasma membrane (**Supplemental Figure 3A,B**). Accumulations of both VEGFR2 and podocalyxin were not evident in the CK-666 treated group as compared with controls; this may be due to the requirement of an established polarity axis for proper trafficking that was absent in 2D culture. We also tested if administration of CK-666 affected CME by live-imaging clathrin puncta before and after drug supplementation. Acute inhibition of Arp2/3 significantly reduced the colocalization of Arf6 with clathrin (**Supplemental Figure 4A,B**). Interestingly, inhibition of CME by addition of Pitstop2 did not affect localization of Arf6 with clathrin (**Supplemental Figure 4C,D**). Overall, these results indicate that global blockade of actin polymerization is capable of inducing protein accumulation similar to loss of Arf6.

### Arf6 is Required for Sprouting Angiogenesis and Lumen Formation

Given depletion of Arf6 was associated with protein sequestration and blunted actin polymerization, we next wanted to determine the requirement of Arf6 for morphogenic behaviors such as sprouting and lumen formation. Arf6 KD produced shorter, thinner sprouts with few discernable lumens (**Figure 5A-E**). There was also an increase in vacuolations (non-contiguous cavities) in Arf6 KD sprouts; this phenotype is a signifier of distorted lumen formation programs[37, 41] (**Supplemental Figure 5A**). In 2D cultured ECs, we did not observe a significant difference in migration via scratch wound assay between Arf6 KD ECs and controls (**Supplemental Figure 5B,C**). These results suggests that loss of Arf6 does not affect migration programs in 2D culture; however, in sprouting scenarios, Arf6 is required for proper sprout formation.

**Figure 5.**
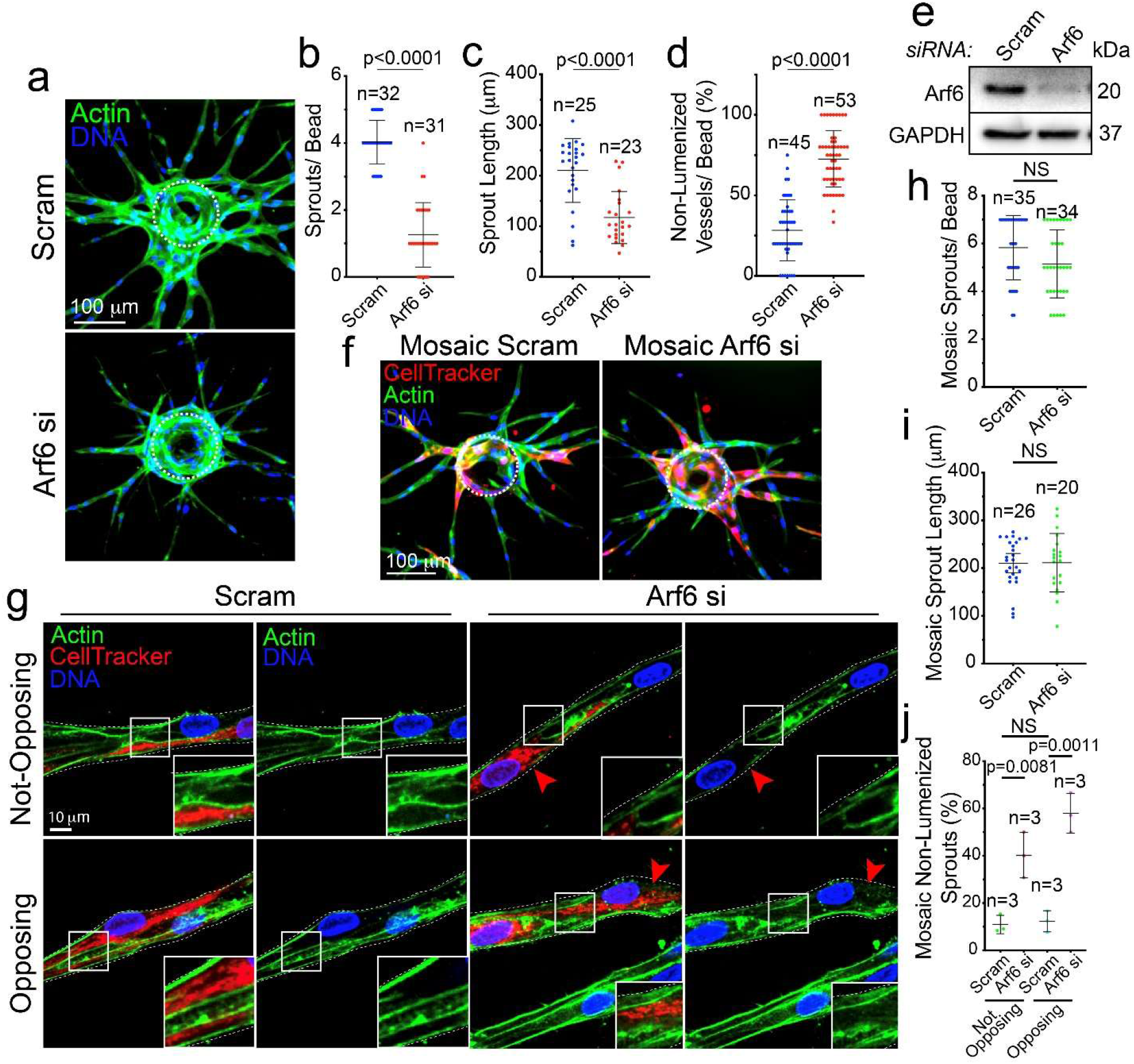
Arf6 is Required for Angiogenic Sprouting. **A.** Representative images of scramble (Scram) control and Arf6 siRNA (si) knockdown (KD) sprouts. **B-D.** Quantification of indicated sprouting parameters between groups. **E.** Western blot confirmation of Arf6 si knock down efficiency. **F.** Representative images of mosaic Scram and Arf6 si sprouts. **G.** Representative images of non-opposing (top panels, an isolated si-treated cell) and opposing (bottom panels, two adjacent si-treated cells) sprout sections. Red arrowheads denote thinned filamentous actin network. **H-J.** Quantification of indicated parameters across groups. In all images L denotes lumen. n = number of sprouts. Error bars represent standard deviation, middle bars are the mean. NS = non-significant. Statistical significance was assessed with an unpaired t-test or a 1-way ANOVA followed by a Dunnett multiple comparisons test. Insets are areas of higher magnification. White dashed lines mark sprout exterior. Dashed circles outline the microbead. All experiments were done using Human umbilical vein endothelial cells in triplicate.

To subvert the global effect of the Arf6 KD on sprouting behaviors, we switched to a mosaic approach. To accomplish this, a population of siRNA-treated ECs were marked with cell tracker, then combined 50:50 with a scrambled-treated control population. Two sprout scenarios were quantified: 1) sprouts with non-opposing KD ECs (KD ECs opposite a WT cell); and 2) sprout areas with two KD ECs opposite each other (opposing). The KD mosaicism rescued sprout length and sprouts per bead to control levels **(Figure 5F-I**). In both scenarios, areas containing individual or opposing Arf6 KD ECs were significantly less lumenized as compared with neighboring WT controls (**Figure 5J**). Notably, KD ECs demonstrated reduced F-actin content as observed previously. Overall, these data suggest Arf6 operates in a cell autonomous fashion and is critical to normal sprout formation during angiogenesis.

### ACAP2 and ARNO Ablation Do Not Phenocopy Arf6 Sprouting Defects

The guanine exchange factor, ARNO and the GTPase activating protein (GAP), ACAP2, have both been reported to regulate cytoskeletal changes through modulation of Arf6 activity[42, 43]. Thus, our next goal was to determine to what extent ARNO and ACAP2 modulated Arf6 function in ECs. Both tagRFP-ACAP2 and GFP-ARNO demonstrated strong colocalization with Arf6 (**Figure 6A,B**). Predictably, loss of ACAP2 increased Arf6 activity, whereas KD of ARNO resulted in reduced Arf6 activation (**Figure 6C-E**). Similar to Arf6 KD, loss of ARNO and ACAP2 resulted in reduced F-actin as compared to controls (**Supplemental Figure 6A,B**). Suppression of ACAP2, and elevated activation of Arf6, did not alter Arf6’s ability to localize to clathrin (**Supplemental Figure 6C,D**). These results support the notion that ACAP2 and ARNO participate in the regulation of Arf6 activity in ECs.

**Figure 6.**
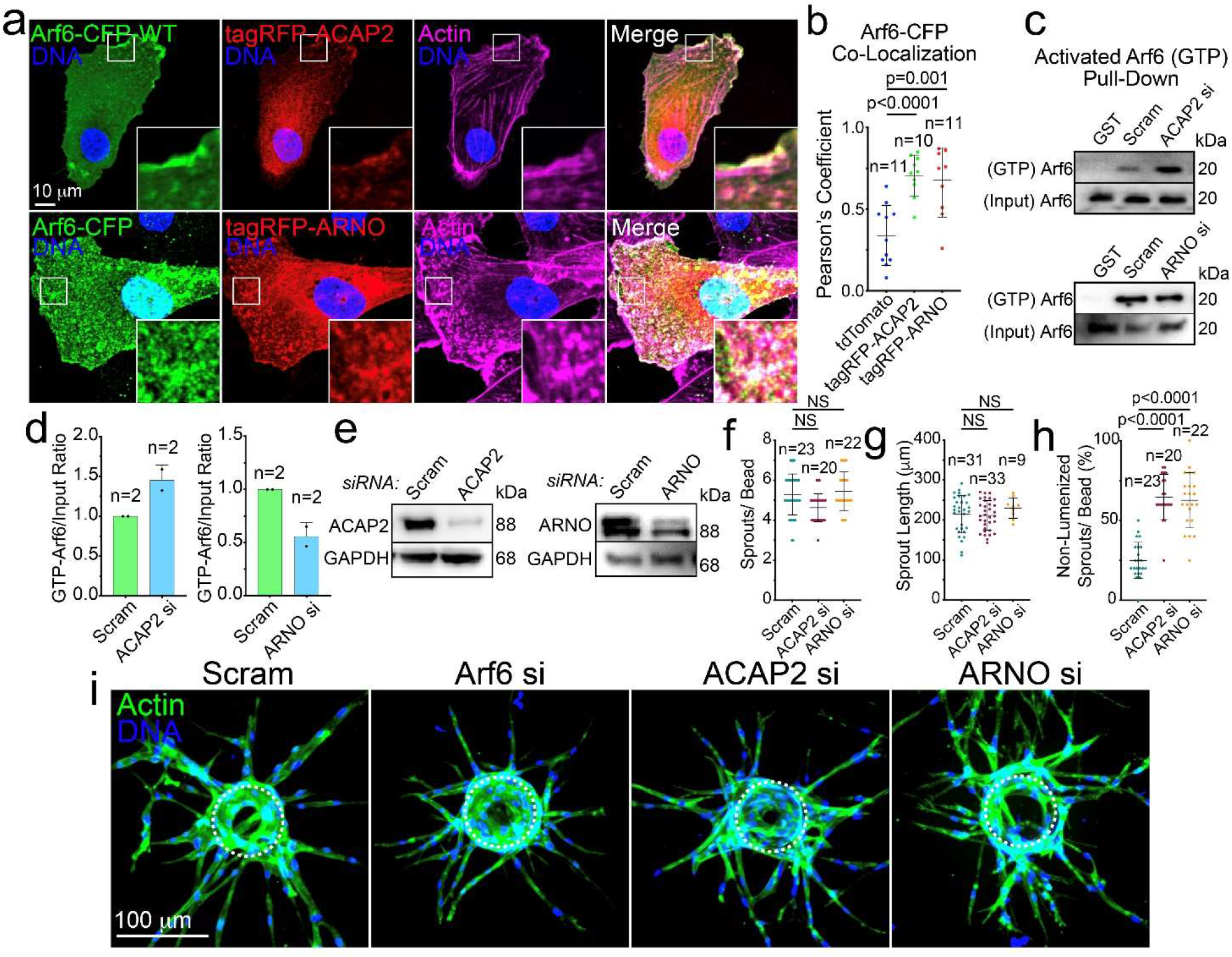
Loss of ACAP2 or ARNO Do Not Phenocopy Loss of Arf6. **A.** Representative images of cells expression wild-type (WT) Arf6-CFP with tagRFP-ACAP2 (top panels) or tagRFP-ARNO (bottom panels). **B.** Pearson’s Coefficient of Arf6-CFP with indicated proteins. n = number of cells. **C.** GTP pulldown assay with GGA3-coated beads to probe for activated Arf6. Cells were treated with scramble (Scram) control, ACAP2 siRNA (si) or ARNO si. **D.** Quantification of band intensity in pull-down blots. n=number of pull-downs. **E.** Western blot confirmation of si knockdown (KD) of ACAP2 and ARNO. **F-H.** Quantification of indicated sprouting parameters between groups. n = number of sprouts. **I.** Representative images of sprout morphology between indicated groups. Dashed circles outline the microbead. In all images white box denotes area of magnification. Error bars represent standard deviation, middle bars are the mean. NS = non-significant. Statistical significance was assessed with an unpaired t-test or a 1-way ANOVA followed by a Dunnett multiple comparisons test. All experiments were done in Human umbilical vein endothelial cells in triplicate.

**Figure 7.**
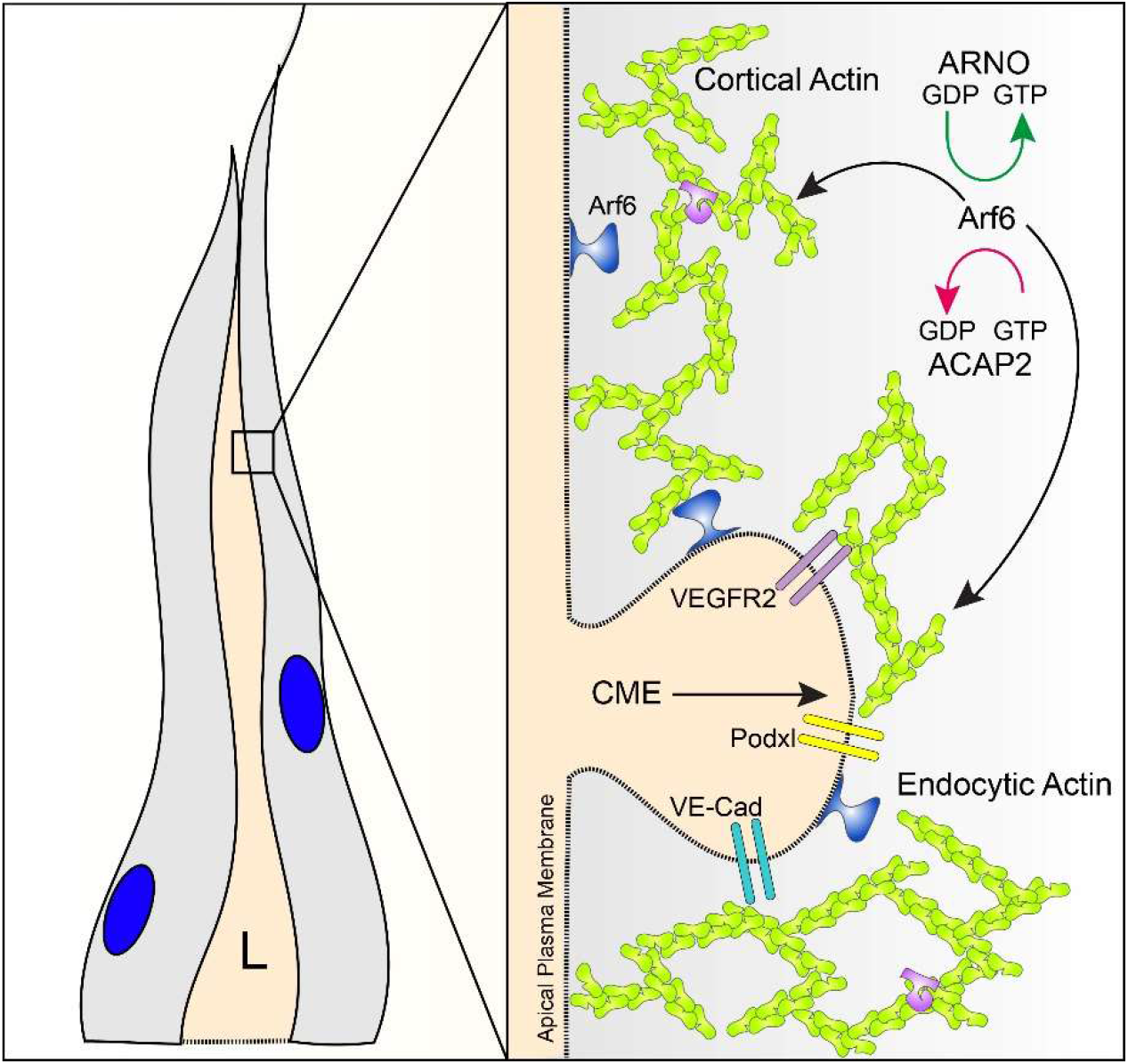
Proposed model for Arf6 Activity in Angiogenesis. In sprouts Arf6 is primarily localized to the apical membrane in close association with the actin cortex and sites of clathrin-mediate endocytosis (CME, membrane invagination). In Arf6’s absence, transmembrane proteins such as vascular-endothelial cadherin (VE-cad), podocalyxin (Podxl) and vascular-endothelial growth factor receptor 2 (VEGFR2) are not correctly internalized. Generally, Arf6 can equally control CME-based and motility-based actin populations through upstream interactions with its guanine exchange factor (GEF) ARNO and GTPase-activating protein (GAP) ACAP2.

Lastly, we determined how loss of ACAP2 or ARNO impacted sprouting behaviors. Unlike Arf6 KDs, ACAP2 and ARNO sprouts showed no significant difference in sprouts per bead or sprout length (**Figure 6F,G**). However, non-lumenized sprouts were significantly higher in both ACAP2 and ARNO KD sprouts (**Figure 6H**). The sprout morphology of ACAP2 and ARNO KDs were similar to Arf6 KD group in their thinner appearance as compared with controls (**Figure 6I**). This data indicates that although ARNO can modulate Arf6 activity, its loss does not completely reprise the Arf6 KD phenotype.

## DISCUSSION

Arf6 has been shown to play an impactful role in blood vessel morphogenesis by way of controlling growth factor signaling capacity. Internalization of trans-membrane proteins, such as receptors, via CME are reliant on Arf6 in providing the actin scaffolding necessary for physical dissociation from the plasma membrane[44]. In endothelial tissue, loss of Arf6 has been leveraged to mitigate chronic growth factor signaling by diminishing CME in diseases such as diabetic retinopathy and solid cancers[33, 34, 45]. However, to date, little has been explored on other potential effects of loss of Arf6 with regard to angiogenic function. This is important as Arf6 and its intimate association with actin regulatory processes may reach far beyond its function in CME. In the current investigation, we took a simple approach in both validating previous Arf6 associations with CME machinery and extended these observations using high-resolution microscopy to uncover how Arf6, and its loss, affected multiple EC behaviors. For the first time, we show where Arf6 localizes in a sprout relative to other apical and basal markers. Additionally, we demonstrate how loss of Arf6 distorts not only the localization of multiple endothelial transmembrane proteins, but its requirement for proper internalization. Our results also highlight the magnitude of influence Arf6 exerts over actin polymerization dynamics, which may explain why Arf6 ablation so dramatically affected angiogenic behaviors. Cumulatively, our results show that Arf6 is not only important for receptor endocytosis but is likely necessary for CME-mediated removal of other critical transmembrane proteins and equally important for modulating non-CME-based actin dynamics.

Our group first became interested in Arf6 due to its reported interaction with the actin modulator Rab35[17]. In exploring Rab35 in ECs, we found that Arf6 was not a critical target of Rab35, but none-the-less, indispensable for proper blood vessel growth. Our initial probe into Arf6 highlighted two major themes: 1) Arf6’s receptor-based interactions were highly characterized in blood vessels; but 2) there was a dearth of information on where Arf6 localized in sprouts as well as its impact on basic angiogenic parameters. This was somewhat surprising given the mounting research inertia on Arf6 as a therapeutic agent. Thus, our goal in the current investigation was to provide an expanded molecular characterization of Arf6 discerning its primary function in angiogenic tissue.

Our first focus was to test how endothelial Arf6 localized to both sites of CME and actin to provide a handle on its primary endothelial function. In testing this, our results demonstrated that Arf6 in ECs equally localize to both structures; thus, doesn’t have a dominant preference. Additionally, Arf6 recruitment was not contingent on having established apicobasal polarity as 2D culture and 3D sprouts demonstrated no difference in colocalization to sites of CME or actin. In multicellular sprouts, Arf6 and clathrin both largely resided at the apical membrane. This finding was rather puzzling as Arf6’s most highly published interactor β1-integrin was localized on the basal surface[39, 46]. Further testing revealed that Arf6 does indeed reduce integrin internalization in agreement with previous reports; however, we could also sequester β1-integrin by inhibiting actin polymerization. This finding supports the notion that Arf6’s actin regulatory function, *per se*, may have a secondary effect in modulating β1-integrin through perturbations in the actin cytoskeleton and downstream mechanotransduction pathways[47, 48].

Uncoupling Arf6’s involvement in generalized CME processes from those specifically targeting cortical actin and cell shape changes would be exceedingly difficult given the overlap in molecular pathways. Despite this caveat, we determined the overall cellular influence Arf6 held on actin polymerization in ECs. Again, we believed this was an important parameter as tissue-wide Arf6 ablation has been successfully performed to combat several vascular diseases. Loss of Arf6 significantly shifted the total cellular actin pool to a predominantly globular state, suggesting a lack of F-actin content. This result is in line with Arf6’s previously established role as a positive regulator of actin polymerization[49–51]. With such a dramatic reduction in actin-related processes, it would be interesting to know how long-term inhibition of Arf6 would impact established blood vessel homeostasis and related cytoskeletal signaling.

A major finding of our study is that Arf6 is required for virtually all aspects of angiogenesis in our model system. Given its importance to actin polymerization this could be predicted; although, to our knowledge, this has not been explicitly tested to date. Loss of Arf6 severely distorted sprout growth characteristics as well as lumen formation parameters. Again, this data further supports the primacy of microfilament regulation in governing normal blood vessel morphodynamics and patterning behaviors[52–55]. This data could be viewed as somewhat paradoxical on the backdrop of several investigations using Arf6 knockout to rectify aspects of vascular dysfunction. Although, vascular restoration was not our primary focus, our results do suggest ablation of Arf6 function can produce ‘collateral damage’ during physiological angiogenesis. This information needs to be fully validated *in vivo*. Nevertheless, taken at face value, these results could be interpreted as cautionary in using Arf6 as a therapeutic target when blood vessels are still undergoing extensive growth, such as in fetal or juvenile development.

Overall, our investigation into Arf6 reinforces its role in CME and furthers the notion that a primary function of Arf6 is controlling actin polymerization in endothelial tissue. Arf6 seems to be equally adept at participating in CME and clathrin-independent processes as well as influencing larger-scale morphodynamic behaviors through regulating cytoskeletal programs; the common denominator being spatiotemporal control of actin dynamics. In endothelial tissue and in vivo blood vessels the Arf6’s cellular interactome is largely unidentified, this is a void in our understanding and only contributes to the opacity of Arf6’s mechanistic reach. To this end, our result in knocking down ARNO demonstrated reduced Arf6 activity, but did not replicate the Arf6 loss of function lumenization defect. Given there are multiple Arf family members with overlapping functions as well as a diverse cadre of GEFs and GAPs, it could be assumed that Arf6 or related members may play a definitive role in many vital biological processes that have yet to be discovered. Indeed, much still needs to be characterized in the way of Arf6 biology to accurately understand its role in blood vessel development, disease progression and therapeutic potential.

**Table 1.** Major Reagents.

| **Reagent** | **Vendor** | **Catalog #** |
| --- | --- | --- |
| OPTI-MEM 1 Reduced Serum Medium, no phenol red | ThermoFisher | 31985070 |
| Polyethyleneamine Branched (PEI) | Sigma-Aldrich | 408727 |
| Chloroquine Diphosphate Crystalline (CQ) | Sigma-Aldrich | C6628-25G |
| Endothelial Cell Growth Medium 2 | PromoCell | C-22011 |
| DMEM, High Glucose, with L-Glutamine | Genesee Scientific | 25-500 |
| GenClone Fetal Bovine Serum (FBS) | Genesee Scientific | 25-514 |
| Penicillin-Streptomycin 100X Solution | Genesee Scientific | P4333-100ML |
| DPBS, no Calcium, no Magnesium | ThermoFisher | 14190250 |
| Trypsin-EDTA, 0.25% 1X, phenol red | Genesee Scientific | 25-510 |
| Paraformaldehyde 20% Aqueous Sol. EM Grade | Electron Microscopy Sciences | 15713 |
| BSA Lyophilized Powder, Fraction V | Genesee Scientific | 25-529 |
| Cytoskeleton G actin/ F actin In Vivo Assay Kit | Cytoskeleton, Inc. | BK037-BK037 |
| Culture-Insert 2 Well in μ-Dish 35 | Ibidi | 81176 |
| Dimethyl Sulfoxide (DMSO) | Sigma-Aldrich | D2650-5X10ML |
| Cytodex Microcarrier Beads | Sigma-Aldrich | C3275-10G |
| Fibrinogen Type 1-S from Bovine Plasma | Sigma-Aldrich | F8630-1G |
| Thrombin from Bovine Plasma | Sigma-Aldrich | T7513-500UN |
| Aprotinin Protease Inhibitor | ThermoFisher | 78432 |
| Phenol-Red (Zebrafish Injection Mixture) | Avantor/ VWR | 34487-61-1 |
| CellTracker Deep Red | ThermoFisher | M22426 |
| BCA Protein Assay Kit | ThermoFisher | 23225 |
| NHLF | Lonza | CC-2512 |
| HEK 293-A | ThermoFisher | R70507 |
| Protease inhibitor cocktail | GoldBio | GB-334-20 |
| Arf6 Pull-Down Activation Assay Biochem Kit | Cytoskeleton, Inc. | BK033 |
| Mem-PER™ Plus Membrane Protein Extraction Kit | ThermoFisher | 89842 |

**Table 2.** Small Molecules.

| Name | Vendor or Source | Catalog No./ Clone | Working Concentration |
| --- | --- | --- | --- |
| PitStop2 | Sigma-Aldrich | SML1169-5MG | 10 uM |
| CK-666 | Sigma-Aldrich | SML0006-5MG | 1uM |

**Table 3.**
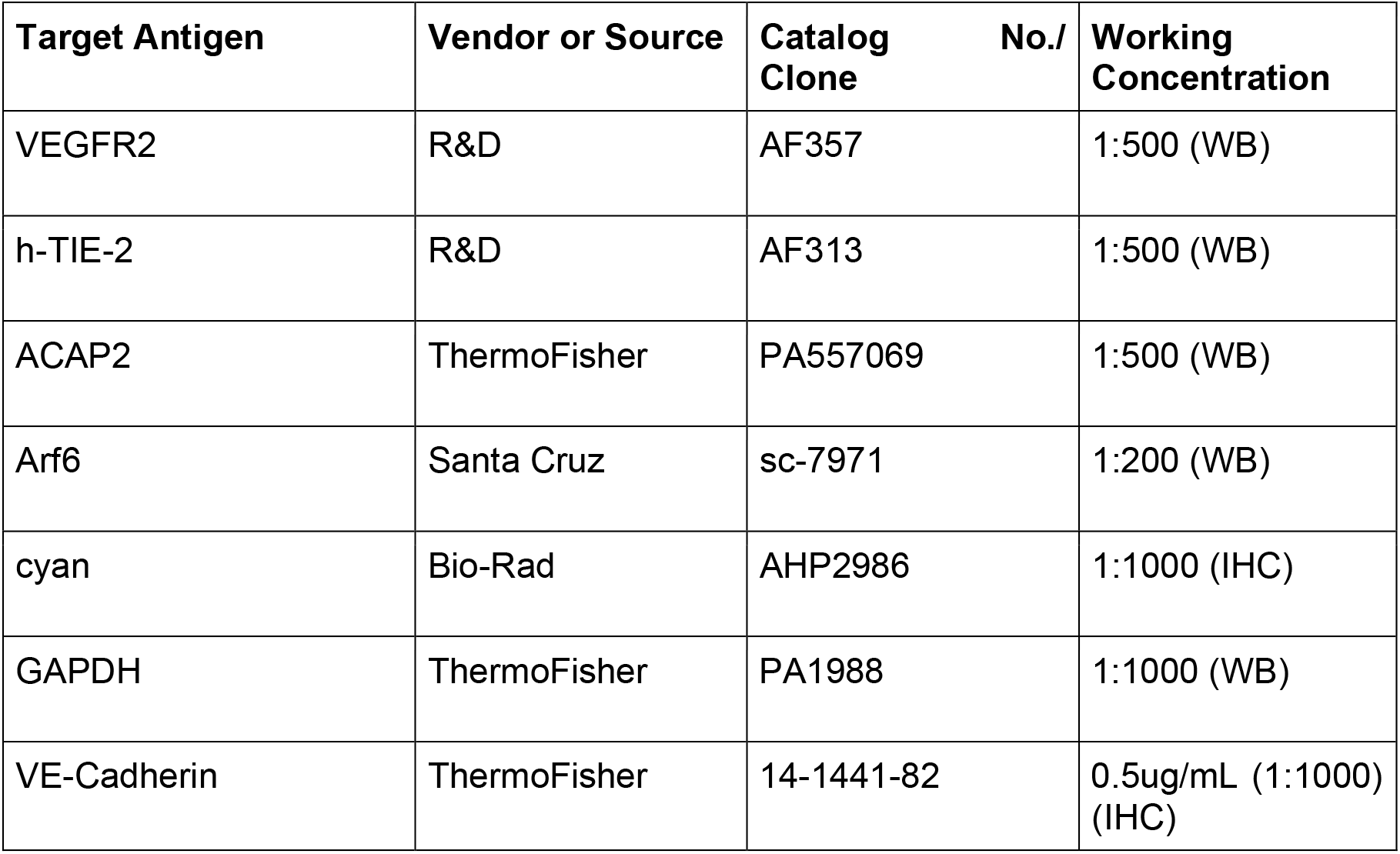
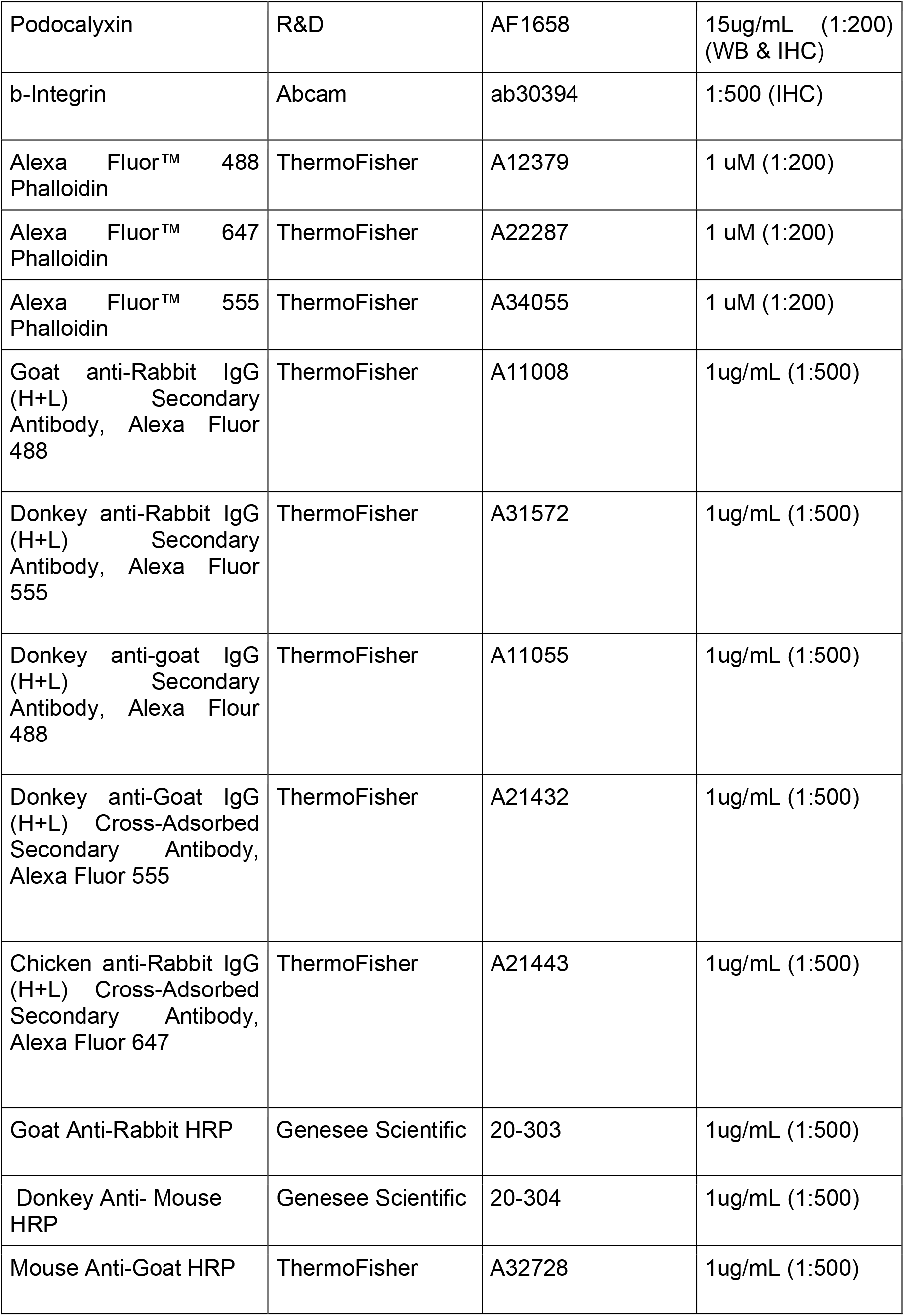
Antibodies.

**Table 4.** siRNA.

| siRNA Target | Vendor | ID # |
| --- | --- | --- |
| Silencer™ Negative Control No. 1 siRNA | ThermoFisher | AM4611 |
| ACAP2 | ThermoFisher | siRNA ID: s24011 |
| ARNO | ThermoFisher | siRNA ID: s225107 |
| Arf6 | ThermoFisher | siRNA ID: s1565 |

**Table 5.** Plasmids.

| Name | Vendor or Source | Catalog No. |
| --- | --- | --- |
| pARF6-CFP | Addgene | 11382 |
| pARF6(T27N)-CFP | Addgene | 11386 |
| pARF6(Q67L)-CFP | Addgene | 11387 |
| mEmerald-ARP2-C-14 | Addgene | 53992 |
| mCherry-ARP2-N-14 | Addgene | 54980 |
| mTagRFP-T-Clathrin-15 | Addgene | 58005 |

## ACKNOWLEDGEMENTS

Work was supported by funding from the National Heart Lung Blood Institute Grant R15HL156106-01A1 (EJK), R01HL155921-01A1) (EJK).

## AUTHOR CONTRIBUTIONS

CRF, MLB and MMS performed all experiments. CRF and EJK planned experiments and wrote the manuscript.

## Supplemental Information

### SUPPLEMENTAL FIGURES

**Supplemental Figure 1.**
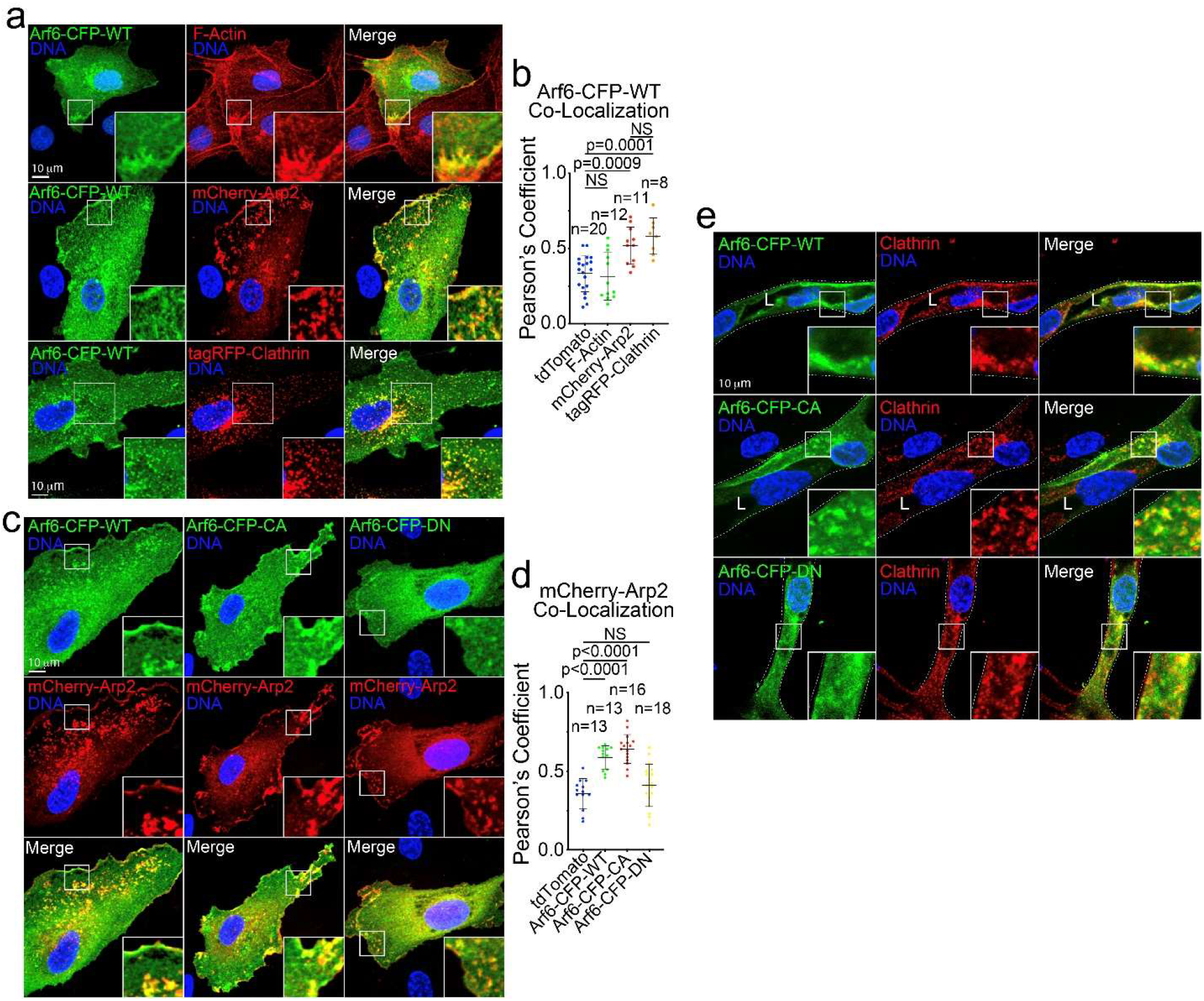
Arf6 Localizes to Cortical Actin and Clathrin. **A**. Representative images of 2-dimensional cells expressing wild-type (WT) Arf6-CFP, mCherry-Arp2, or tagRFP-Clathrin. **B**. Pearson’s coefficient of Arf6-CFP with indicated proteins. **C**. Representative images of WT, constitutively-active (CA), or dominant-negative (DN) Arf6-CFP with mCherry-Arp2. **D**. Pearson’s coefficient of mCherry-Arp2 with indicated proteins. **E**. Representative images of WT, CA, DN Arf6-CFP stained for endogenous clathrin. N = number of cells. NS=non-significant. Error bars represent standard deviation, middle bars are the mean. Statistical significance was assessed with an unpaired t-test or a 1-way ANOVA followed by a Dunnett multiple comparisons test. All experiments were performed using human umbilical vein endothelial cells in triplicate.

**Supplemental Figure 2.**
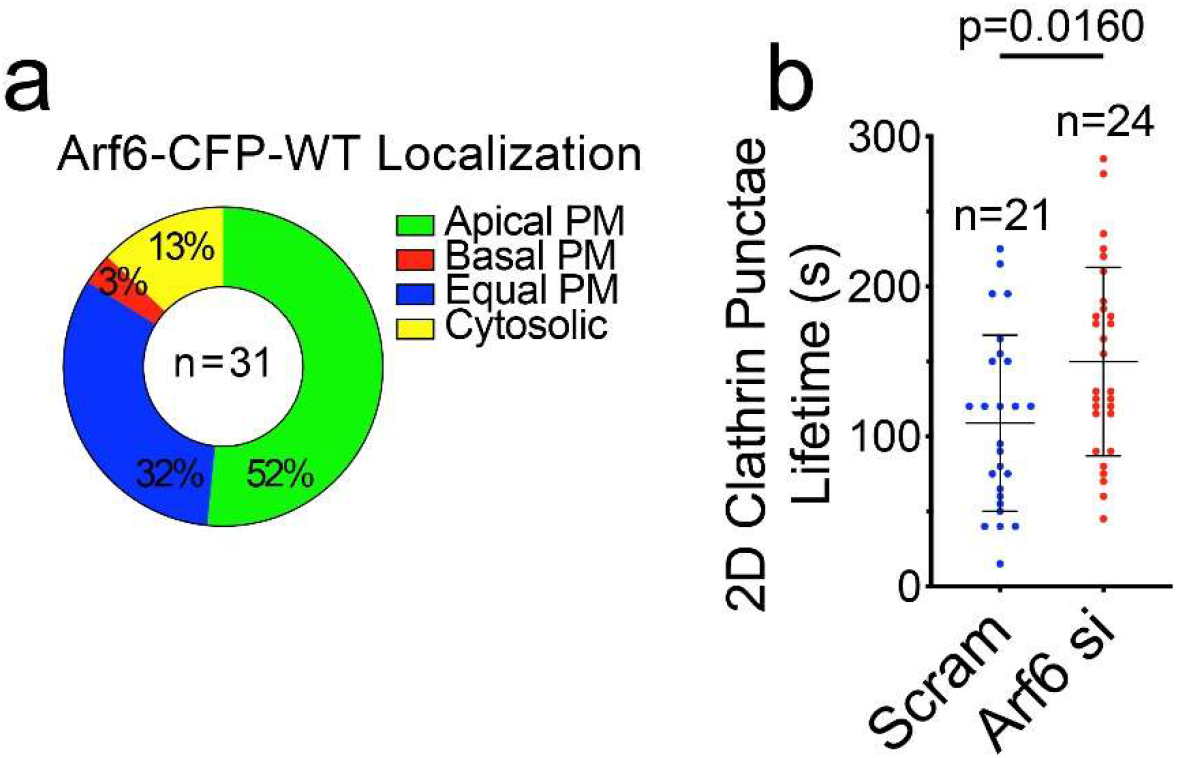
Arf6 Localization Preference and Impact on Clathrin Turnover. **A**. Quantification of wild-type (WT) Arf6-CFP cellular localization preference to the apical membrane (PM), basal membrane, equally localized to the basal and apical membrane, or localized primarily to the cytoplasm. **B**. Quantification of clathrin puncta lifetime in scramble (Scram) and Arf6 siRNA knockdown cells. N = number of cells. Error bars represent standard deviation, middle bars are the mean. Statistical significance was assessed with an unpaired t-test or a 1-way ANOVA followed by a Dunnett multiple comparisons test. All experiments were done using human umbilical vein endothelial cells in triplicate.

**Supplemental Figure 3.**
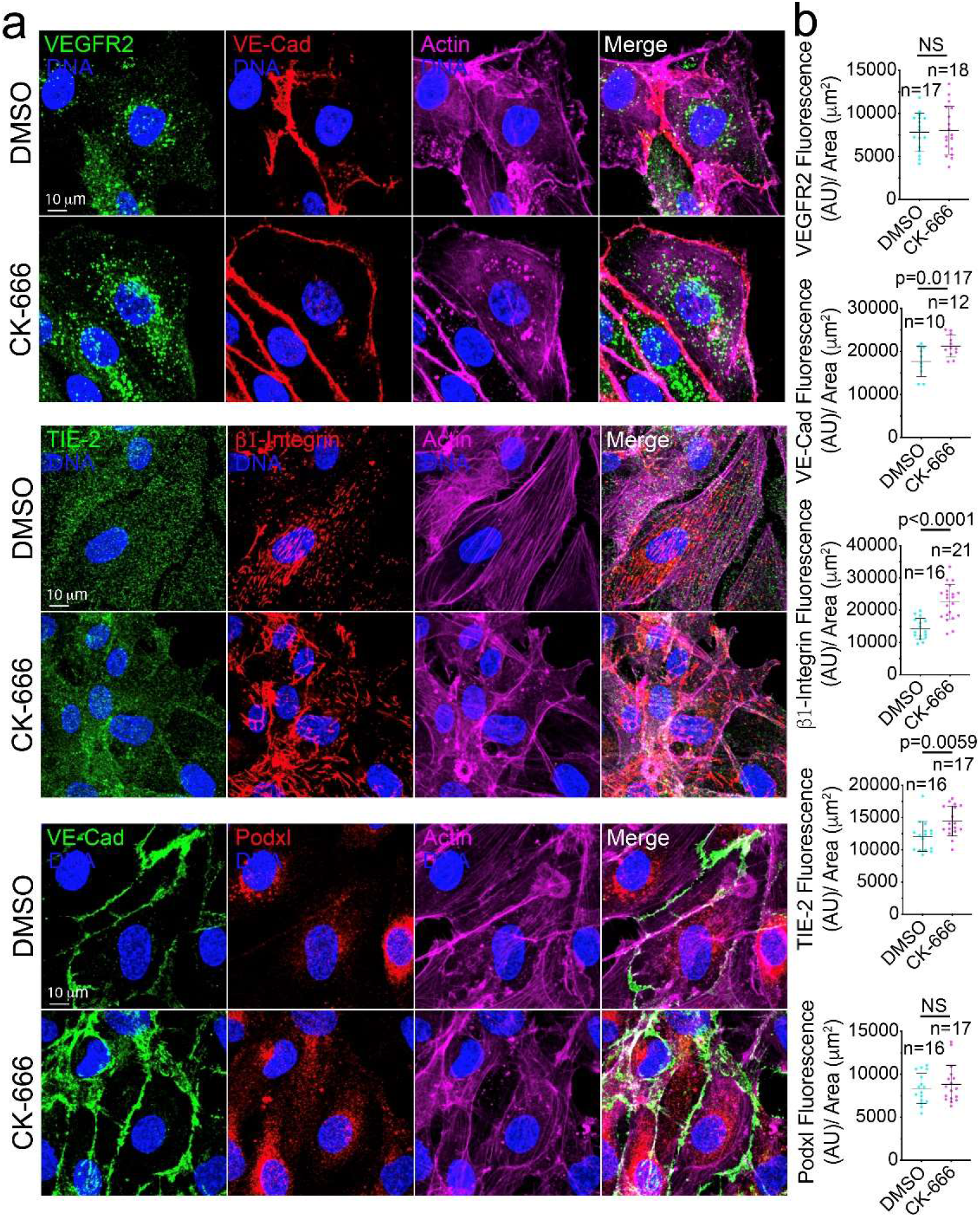
Actin Polymerization is Required for Endocytosis. **A**. Representative images of cells treated with DMSO (control) or Arp2/3-inhibitor (CK-666) and stained for indicated proteins. White arrows are indicative of direction of cell migration. **B**. Quantification of fluorescent intensity of indicated proteins normalized to cell area. Error bars represent standard deviation, middle bars are the mean. Statistical significance was assessed with an unpaired t-test or a 1-way ANOVA followed by a Dunnett multiple comparisons test. N = number of cells. NS=non-significant. All experiments were done using human umbilical vein endothelial cells in triplicate.

**Supplemental Figure 4.**
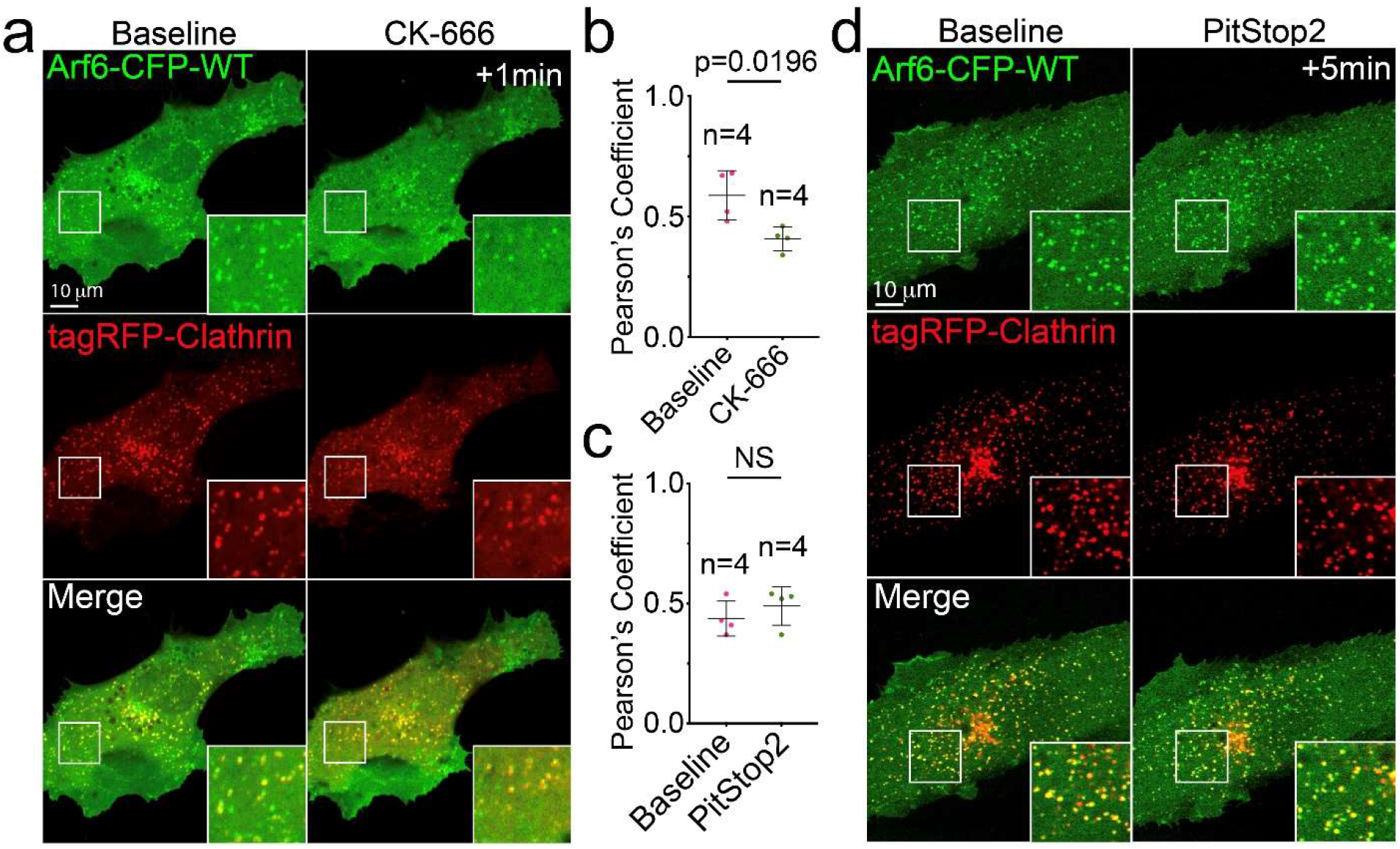
Cortical Actin Improves Arf6 and Clathrin Colocalization. **A.** Live imaging of wild-type (WT) Arf6-CFP with tagRFP-Clathrin at baseline and after treatment with CK-666. **B**. Pearson’s Coefficient of WT Arf6-CFP and tagRFP-Clathrin between DMSO and CK-666 treated. **C**. Pearson’s Coefficient of WT Arf6-CFP and tagRFP-Clathrin between DMSO and Pitstop treated cells. **D**. Live imaging of WT Arf6-CFP with tagRFP-Clathrin at baseline and after addition of PitStop2. In all images white box denotes area of magnification. Error bars represent standard deviation, middle bars are the mean. Statistical significance was assessed with an unpaired t-test. N = number of cells. NS = non-significant. All experiments were done in Human umbilical vein endothelial cells in triplicate.

**Supplemental Figure 5.**
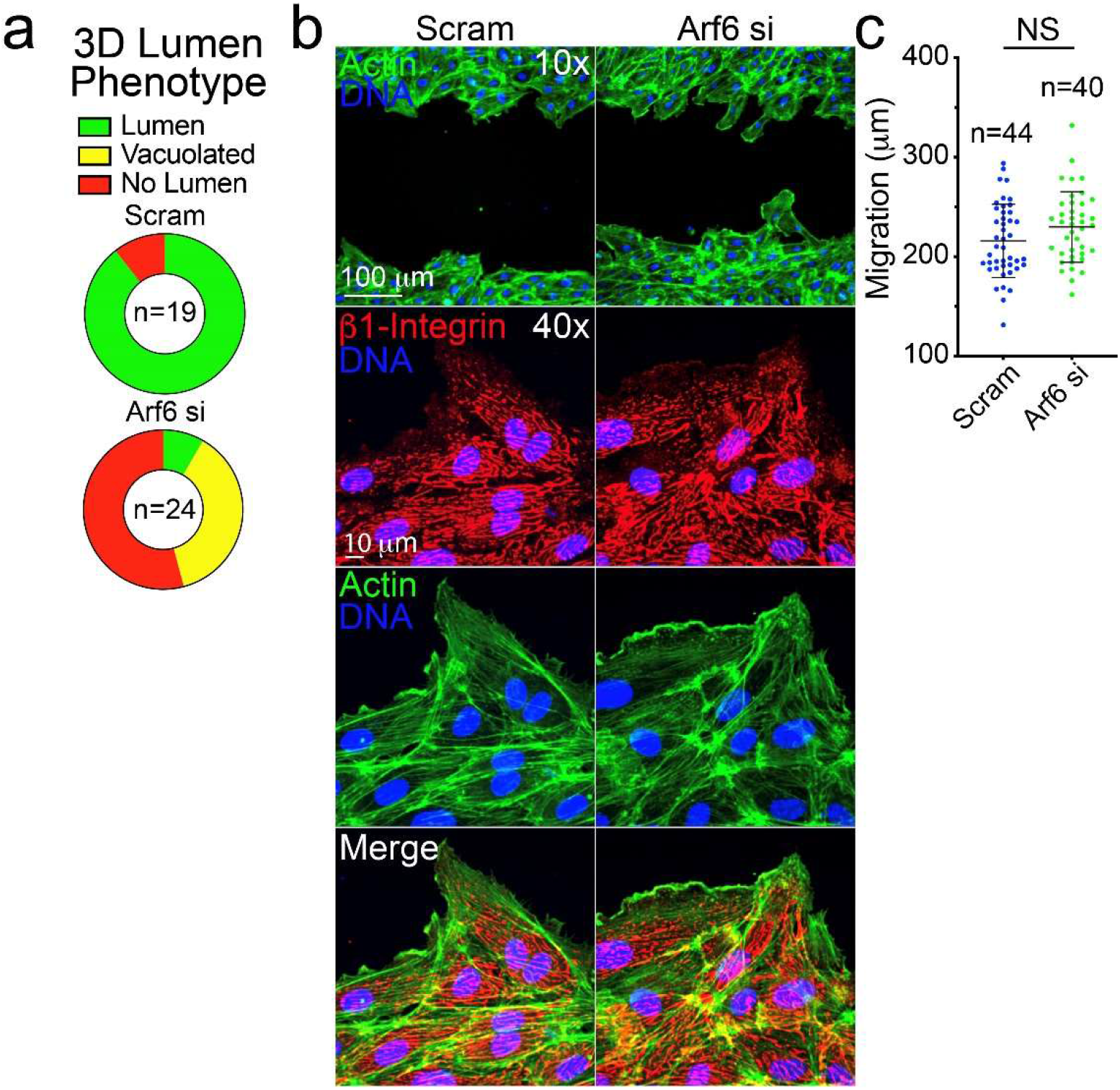
Loss of Arf6 Results in Lumen Defects, but Does Not Impact Cell Motility. **A**. Quantification for lumen phenotypes in scramble (Scram) control and Arf6 siRNA knockdown (KD) sprouts. Vacuolated was defined as large round vacuoles with no contiguous lumen. **B**. Scratch wound assay in which cells were treated with Scram or Arf6 siRNA (si). Cells were stained for β1-integrin and actin. **C**. Quantification migration distance. N=number of measurements. Error bars represent standard deviation, middle bars are the mean. Statistical significance was assessed with an unpaired t-test. All experiments were done using human umbilical vein endothelial cells in triplicate.

**Supplemental Figure 6.**
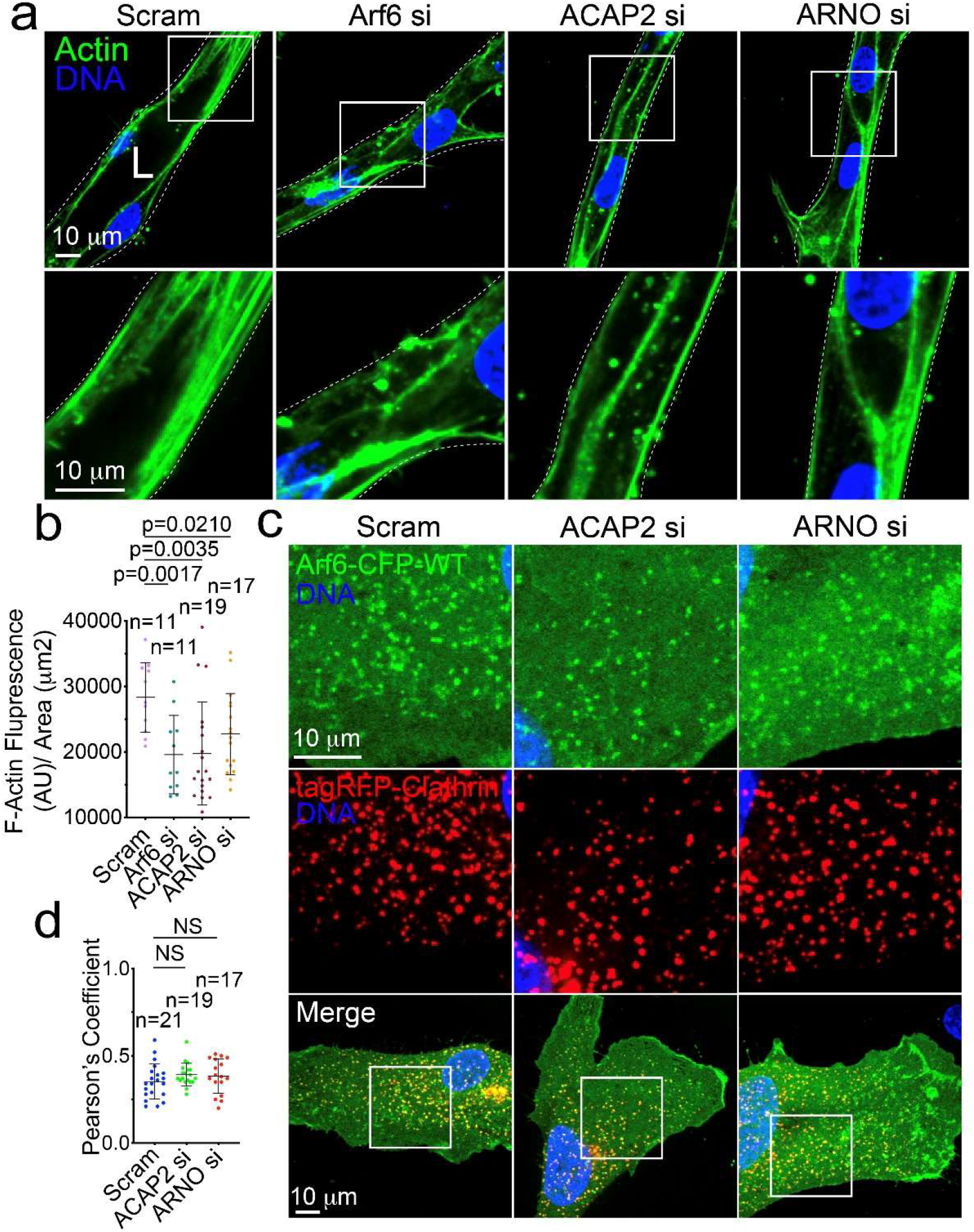
Arf6 Influences Actin Content and Impact of ARNO and ACAP2 on Clathrin Recruitment. **A**. Representative sprout stained for actin with and without Arf6 siRNA (si) knockdown. **B**. Quantification of actin intensity between indicated groups. N=number of sprouts. **C**. Representative images of Wild-type (WT) CFP-Arf6 and tagRFP-Clathrin expressing cell with indicated si-treatment. **D**. Pearson’s Coefficient of CFP-Arf6 and tagRFP-Clathrin between indicated groups. N=number of measurements. Error bars represent standard deviation, middle bars are the mean. Statistical significance was assessed with an unpaired t-test. All experiments were done using human umbilical vein endothelial cells in triplicate.

